# IsRNAcirc: 3D structure prediction of circular RNAs based on coarse-grained molecular dynamics simulation

**DOI:** 10.1101/2024.07.03.601849

**Authors:** Haolin Jiang, Yulian Xu, Yunguang Tong, Dong Zhang, Ruhong Zhou

## Abstract

As an emerging class of RNA molecules, circular RNAs play pivotal roles in various biological processes, thereby determining their three-dimensional (3D) structure is crucial for a deep understanding of their biological significances. Similar to linear RNAs, the development of computational methods for circular RNA 3D structure prediction is challenging, especially considering the inherent flexibility and potentially long length of circular RNAs. Here, we introduce an extension of our previous IsRNA2 model, named IsRNAcirc, to enable circular RNA 3D structure predictions through coarse-grained molecular dynamics simulations. The workflow of IsRNAcirc consists of four main steps, including input preparation, end closure, structure prediction, and model refinement. Our results demonstrate that IsRNAcirc can provide reasonable 3D structure predictions for circular RNAs, which significantly reduce the locally irrational elements contained in the initial input. Moreover, for a validation test set comprising 34 circular RNAs, our IsRNAcirc can generate 3D models with better scores than the template-based 3dRNA method. These findings demonstrate that our IsRNAcirc method is a promising tool to explore the structural details along with intricate interactions of circular RNAs.

**Author Summary:** Knowledge of the 3D structure of circular RNA molecules can help us better understand their roles in eukaryotic cells. However, experimental determination of circular RNA 3D structure remains challenging, with currently only one complete circular RNA 3D structure available in the Protein Data Bank. Thus, extending from their linear counterparts, computational methods that can predict circular RNA 3D structure are urgently needed. In particular, physics-based *de novo* 3D structure prediction methods are promising due to their powerful sampling capability in the conformational space, which avoid the problem of limited available templates in template-based approaches. We have developed IsRNAcirc, a new physics-based method that employs an accurate coarse-grained force field and replica-exchange molecular dynamics simulations to predict the 3D structure of circular RNAs. IsRNAcirc is shown to significantly reduce the number of locally irrational structures in the input trials, which are caused by the lack of suitable templates and the improper assembly of fragments. Furthermore, in the absence of experimental reference structures, the predicted 3D models were validated using the popular scoring functions, such as DFIRE-RNA and rsRNASP, demonstrating that IsRNAcirc outperforms the template-based 3dRNA method in generating low-energy conformations.

## Introduction

Circular RNA molecules are a unique class of RNA that do not have a cap structure at the 5’ end and a poly(A) tail at the 3’ end, making them different from traditional linear RNAs. These circular RNAs are formed through a process called back splicing, in which segments of mRNA precursors are covalently joined together to form a circular structure. The initial and terminal junctures within a circular RNA sequence, known as the circularization sites, are specifically referred to as back-splicing junction (BSJ) regions. The discovery of circular RNAs dates back more than four decades, when the first circular RNA molecule was identified in plant viroids[1]. To date, thanks to advances in high-throughput RNA sequencing (RNA-seq) and specialized bioinformatics tools, thousands of circular RNAs have been discovered in various eukaryotic organisms, including fungi, protists, plants, worms, fish, insects, mammals, etc.[2–5]. Interestingly, many of these circular RNAs originate from protein-coding genes, suggesting that they participate in diverse biological processes and perform multiple functions within cells. For example, a prominent function of circular RNAs is their ability to act as microRNA sponges, sequestering miRNAs and preventing them from targeting and degrading specific mRNAs[6,7]. Other discovered functions of circular RNAs include interacting with ribosomes to increase protein synthesis, interacting with RNA-binding proteins (RBPs) to modulate their activity[8,9], directing proteins to specific cellular locations[2], and translating into unique peptides *via* internal ribosome entry sites (IRES)[10–13]. These rich functions of circular RNAs suggest that they should adopt specific three-dimensional (3D) structures.

However, experimental determination of circular RNA 3D structures remains very challenging, mainly due to the inherent flexibility and potentially considerable length of these RNA molecules. As a result, there is currently only one complete circular RNA 3D structure (PDB ID: 2OIU[14]) deposited in the Protein Data Bank (PDB). Therefore, to facilitate our understanding of the mechanisms of their biological functions, developing computational methods to predict the 3D structure of circular RNAs is a much-needed but lacking task. For their linear counterparts, various computational approaches have been developed over the past few decades to predict RNA 3D structure. In general, these computational approaches can be divided into two different categories: template-based methods and *ab initio* predictions. Template-based methods, such as ModeRNA[15], Vfold[16,17], RNAcomposer[18], and 3dRNA[19,20], rely on known RNA structures of similar sequences as templates to predict structures. The main bottleneck of template-based methods is that the available templates deposited in the PDB are limited, so they can easily fail to provide reasonable predictions for unknown RNA sequences. On the other hand, *ab initio* prediction methods, including NAST[21], iFoldRNA[22], SimRNA[23], and our IsRNA[24–26], utilize molecular dynamics (MD) or Monte Carlo simulations to search the conformational space and a particular scoring scheme (usually the energy functions) to select the lowest energy or most probable structure from the conformation ensemble as the prediction(s). Due to their template-free nature, *ab initio* methods can in principle provide predictions for any RNA sequences, but at a relatively high computational cost. Recently, artificial intelligence (AI)-based approaches have also exhibited great promise in predicting linear RNA 3D structures[27–29]. Nevertheless, coming back to circular RNAs, there is currently only one computational method that can predict their 3D structures. That is, Xiao and coworkers[19] extended their 3dRNA model to enable the prediction of the 3D structures of circular RNAs. Considering the template-based nature of the 3dRNA model and the limited number of available templates, it is necessary to develop other computational methods to predict circular RNA structures, such as *ab initio* prediction methods.

Here, we introduced an extension of our previous IsRNA2 model, termed IsRNAcirc, to enable circular RNA 3D structure predictions *via* coarse-grained (CG) MD simulations. To evaluate the performance of IsRNAcirc, we collected a test set consisting of 34 circular RNAs of different lengths and topologies from the previous study[30]. The results demonstrate that IsRNAcirc can provide reasonable 3D structure predictions for circular RNAs and significantly reduce the locally irrational elements contained in the initial input. Furthermore, since the true structures are unavailable, a comparison of IsRNAcirc and 3dRNA was conducted by model assessments with two knowledge-based scoring functions. Notably, IsRNAcirc outperforms 3dRNA because the former generates conformations with lower energy in different scoring schemes. The presented IsRNAcirc model provides a feasible approach to capture the intricate features of circular RNA structures and facilitates our understanding of their various functions.

## Results and Discussions

### Overview of IsRNA2 coarse-grained model

To predict the 3D structures of linear RNAs, IsRNA2[26] incorporates three essential features (Fig 1A): a five-bead per nucleotide CG representation to preserve the three interacting edges of nucleobases, an accurate CG force field derived from the iterative simulated reference state approach, and the utilization of replica-exchange molecular dynamics (REMD) simulations to effectively sample the conformational space. Using RNA sequence, secondary (2D) structure, and initial 3D structures (optionally) as input, the workflow of IsRNA2 contains sampling the conformational space through REMD simulations with 10 replicas possessing different temperatures, constructing a conformational ensemble (containing 5,000 structure snapshots) from simulation trajectories, clustering the top 10% structures with the lowest potential energies to generate the most probable predictions, and finally recovering all-atom structures and energy minimization (Fig 1B). Our benchmark test showed that IsRNA2 achieves comparable performance to the atomic model in *de novo* modeling of noncanonical RNA motifs and can refine large RNA 3D models predicted by other programs. More details on IsRNA2 can be found in our previous work[26].

**Figure 1.**
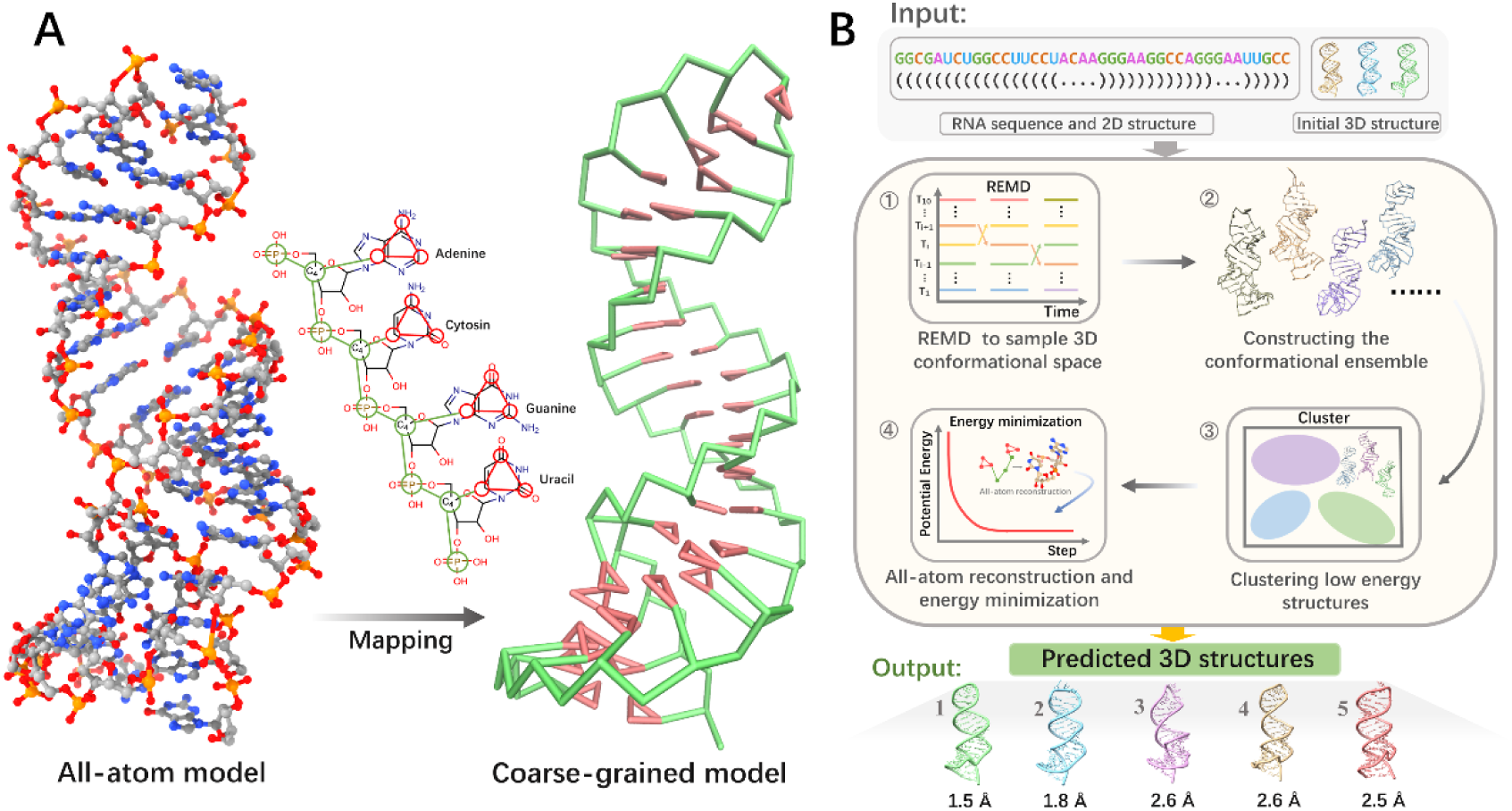
Overview of the coarse-grained (CG) IsRNA2 model for linear RNA 3D structure prediction. (A) Converting an all-atom model (PDB ID:1Z5C, left) into the CG representation (right) through predefined mapping relationships. In IsRNA2 model, the backbone (colored by green) is represented by two CG beads (bead P located on the atom P for the phosphate group and bead S located on the atom C4’ for the ribose ring) and the nucleobase (colored by pink) is represented by three CG beads (each bead located at the center of mass of the related heavy-atom group). (B) An illustrated example to display the workflow for linear RNA 3D structure prediction by IsRNA2 model. Using sequence, 2D structure, and initial 3D structure as input, IsRNA2 employs a four-step process (conformation sampling through REMD, conformational ensemble construction, clustering procedure, and all-atom reconstruction and energy minimization) to generate the predicted 3D structures. For reference, the RMSD of each predicted model relative to the native structure (PDB ID: 1Z5C) was also shown.

### The framework of IsRNAcirc for circular RNA 3D structure prediction

IsRNAcirc is an extension of our previous CG IsRNA2[26] model to enable prediction of circular RNA 3D structure. The main difference in circular RNAs compared to linear RNAs is the presence of covalently bound ends, such that the connection between the two "end nucleotides" (including bonds, bond angles, and torsion angles) is the same as the connection between any two adjacent nucleotides in the sequence. Thus, circular RNAs may adopt different 2D and 3D structures compared to their linear counterparts. There are four main steps to predict circular RNA 3D structure using IsRNAcirc model, including input preparation, end closure, structure prediction, and model refinement (Fig 2). Firstly, from the sequence information, cRNAsp12 tool[31] was used to predict the 2D structure of circular RNA molecules. And based on the sequence information and predicted 2D structure, RNAComposer[18] was utilized to generate three (if available) initial 3D structures for the corresponding linear counterpart. Collectively, the sequence information, the predicted 2D structure, and those obtained initial 3D structures serve as input to IsRNAcirc (Fig 2A). Secondly, in the end closure step, we combined a harmonic potential constraint with a gradually increased force constant and the simulated annealing simulation technique to close the 5’- and 3’-end of the initial 3D structures (Fig 2B). Meanwhile, in order to reduce unreasonable topologies caused by improper fragment assembly in the initial 3D structures, the base-pairing interactions were also weakened in the early stages of simulated annealing simulations. The most probable conformation (obtained by clustering analysis) in the later stage of the simulation was selected as the starting structure for subsequent simulations. This end closure step was specifically designed for circular RNA 3D structure prediction, which is the primary difference between IsRNAcirc and the original IsRNA2 model. Thirdly, once the two “terminal nucleotides” were circularized, standard bonded energy parameters (including bond, bond angles, and torsion angles) were introduced to describe their connection. And a similar process to IsRNA2 was performed to predict circular RNA 3D structures. That is, sampling the conformational space via REMD simulation, constructing the conformation ensemble and clustering the top 10% lowest energy structures, and all-atom reconstruction (Figs 1B and 2C). Considering the longer size of circular RNAs, the stimulation time used in IsRNAcirc (100 ns) is longer than the original IsRNA2 model (50 ns). We chose the top five predicted models (centroid structures of the top five largest clusters) after all-atom recovery as candidate 3D structures of certain circular RNA. Finally, since the CG nature of IsRNAcirc may lead to the loss of fine structural details, a model refinement step was performed based on QRNAS[32] (Fig 2D), which can effectively eliminate the imperfect stereochemical features, unnatural bond lengths, and spatial steric resistance present in the predicted 3D models. Overall, through the above four steps, users can use IsRNAcirc to generate five predicted 3D structure models of circular RNAs in PDB format.

**Figure 2.**
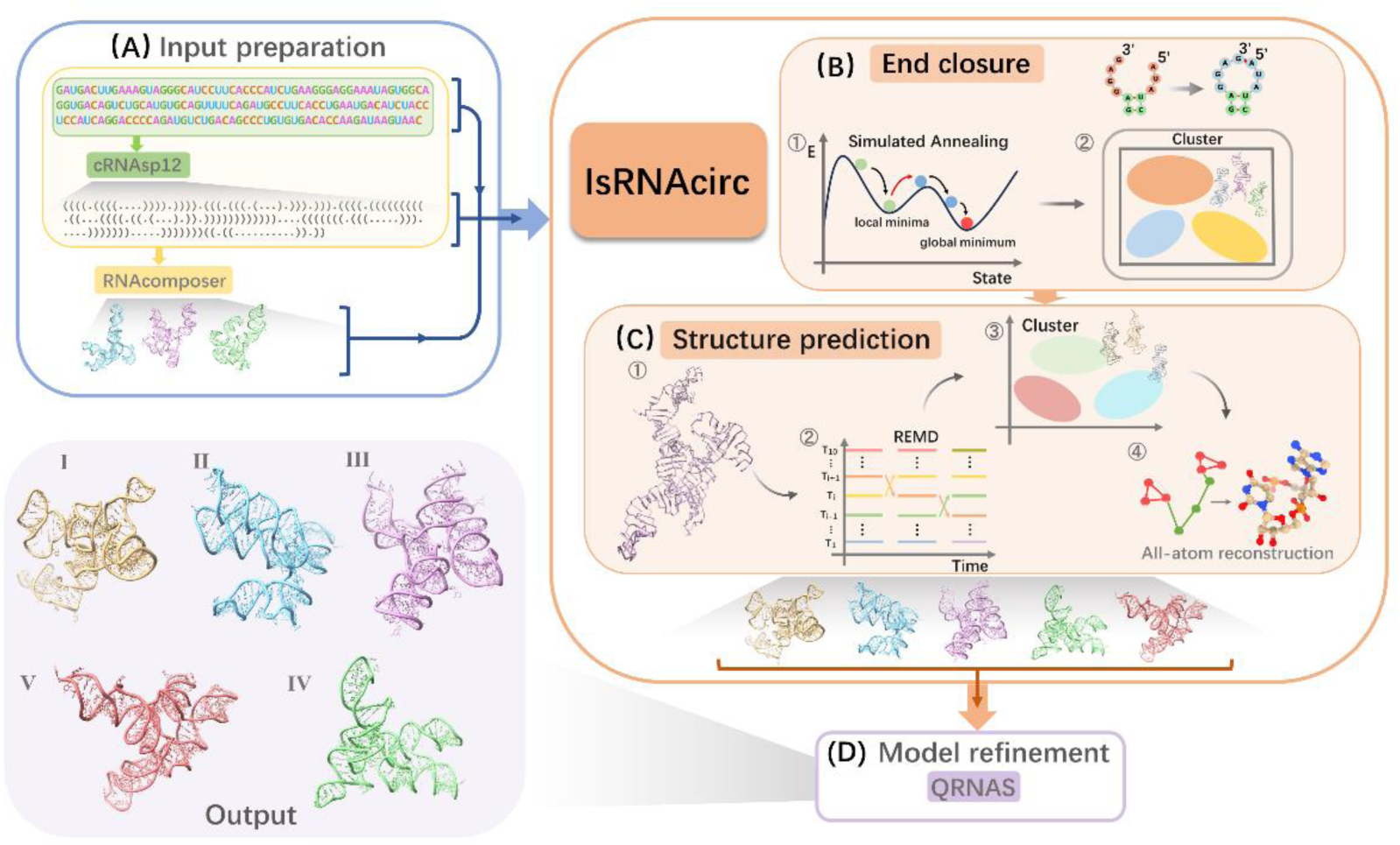
A framework for predicting circular RNA 3D structures using IsRNAcirc. (A) Input preparation. From sequence information, cRNAsp12[31] was used to predict the 2D structure of circular RNAs. Based on the sequence and predicted 2D structure, RNAComposer[18] was used to generate initial 3D structures of the corresponding linear counterpart. Sequence information, predicted 2D structure, and those obtained initial 3D structures are used as inputs for IsRNAcirc for circular RNA 3D structure predictions. (B) End closure. The 5’- and 3’-end of circular RNA are circularized using a harmonic restraint between the two ends of linear structure and the simulated annealing technique. A clustering procedure was performed based on the last 10% MD trajectories to obtain starting structures for the subsequent step. (C) Structure prediction. Possible 3D structures of circular RNA were predicted following a similar process to IsRNA2, including conformational sampling through REMD simulations, clustering the lowest energy conformations to generate the most probable structures, and all-atom reconstruction. By default, five predicted 3D models are provided. (D) Model refinement. QRNAS[32] was employed to refine the predicted 3D models by eliminating problems such as spatial steric resistance and bond breakage.

### Benchmarking of IsRNAcirc on 34 circular RNAs

The predictive ability of IsRNAcirc was tested on a dataset from Chen’s group[30]. This dataset comprises 34 circular RNAs with lengths ranging from 161 to 435 nucleotides. Similar to the previous study[19], those 34 circular RNAs were classified into four types according to the location of BSJ in the 2D structure: helix-circular, hairpin-circular, internal-circular, and junction-circular RNAs. For each type of circular RNA, the two “terminal nucleotides” are well circularized in the 3D models predicted by IsRNAcirc (Fig S1). Moreover, we aligned the predicted circular RNA 3D models with the initial 3D structures of the corresponding linear RNAs (obtained by RNAComposer) and calculated their root mean square deviations (RMSDs). For all four types of circular RNAs, the predicted 3D models differ significantly from the initial linear 3D structures, with average RMSD values larger than 30 Å (Fig S2). This result is different from observations in 3dRNA[19], which showed that helix-circular and hairpin-circular RNAs adopt almost the same folded structures as their linear counterparts, whereas the assembled structures of internal-circular and junction-circular RNAs are very different from those of their linear counterparts. Nevertheless, given the much stronger sampling capacity of IsRNAcirc in the conformational space relative to templated-based approaches (such as 3dRNA and RNAComposer), we believe that our results (large differences between the 3D structures of all circular RNAs and their linear counterparts) are acceptable. Furthermore, a positive correlation was observed between the lengths of circular RNAs and the calculated RMSD values, which implies that longer circular RNAs tend to adopt more different folded conformations relative to the initial linear 3D structures. This point further emphasized the stronger conformational sampling capability of our IsRNAcicr model.

In particular, through view inspection, we observed a lot of locally irrational topologies in the initial 3D structures generated by RNAComposer. Specifically, four main types of locally irrational topologies were identified, including template-free loop, backbone tangle, base collision, and sharp twist. Figs 3A-3D display illustrated examples for each of these four types of locally irrational topologies. Due to the template-based nature of RNAComposer approach and the long length (> 160 nucleotides) of RNA molecules, these locally irrational topologies are thought to be mainly caused by the lack of appropriate templates in the library and improper assembly of the template fragments. For instance, the most prevalent type of locally irrational topology is base collision, in which a backbone segment traverses through the space between two adjacent nucleobases and causes severe atom clashes (Fig 3C). Failure to consider possible overlaps between different template fragments during assembly process is thought to be responsible for the presence of bases collision. On the other hand, because of the lack of template for long loops (≥15 nucleotides in length) in the library, an alternative single strand with predefined geometries was introduced in the initial 3D structures, which results in the second most common type of locally irrational topology (termed template-free loop, Fig 3A).

**Figure 3.**
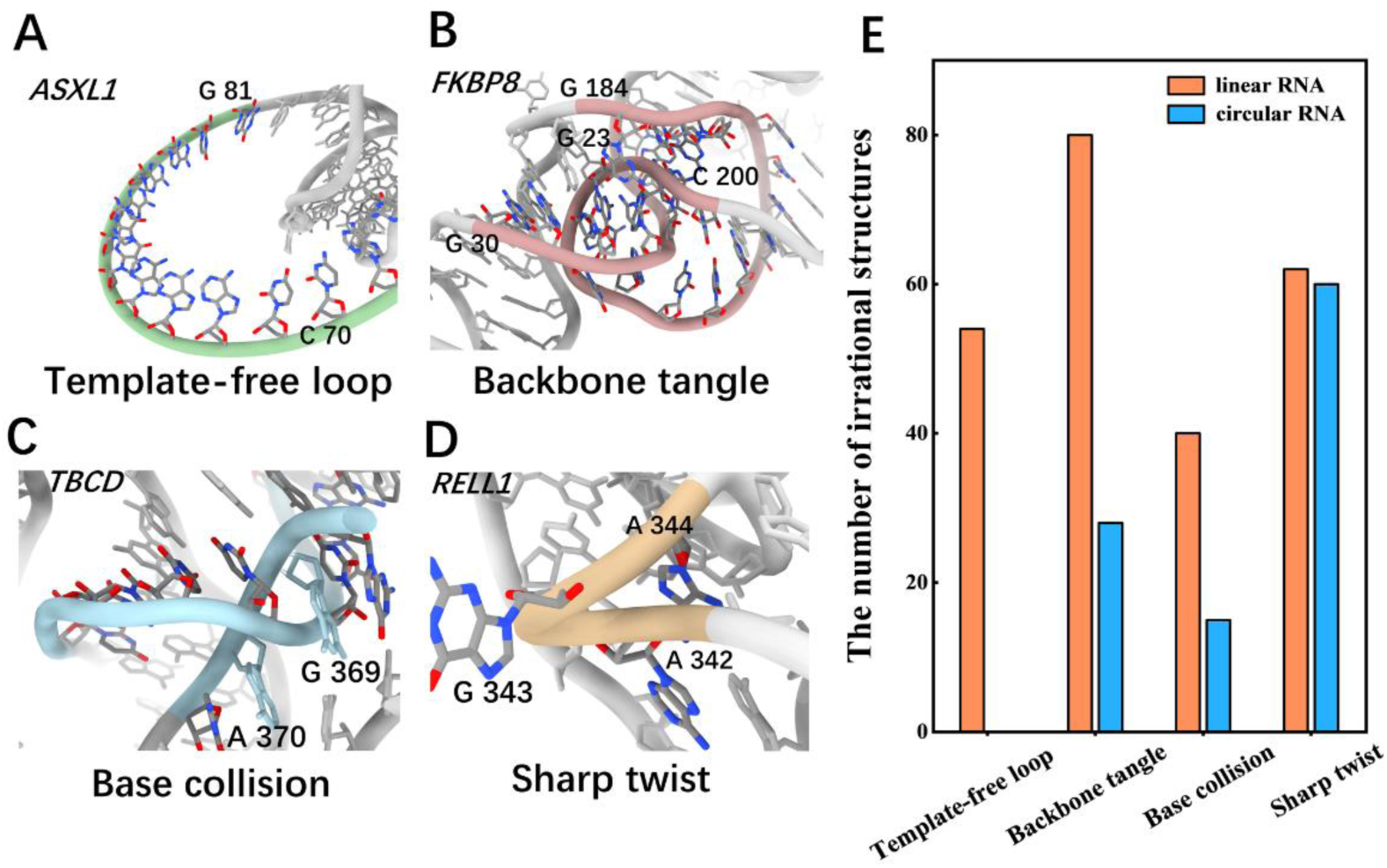
IsRNAcirc can significantly reduce the locally irrational topologies contained in initial 3D structures (generated by RNAComposer). (A-D) Four types of locally irrational topology were identified in initial 3D structures through view inspection: (A) template-free loop due to a lack of related templates in the library, (B) backbone tangle caused by improper template assembly, (C) base collision, in which a backbone segment passes through the middle of two nucleobases, and (D) sharp twist, in which an apparently unreasonable sharp turn is observed in the backbone. The names of circular RNAs for which the relevant irrational topology was observed in the initial 3D structures were also given in top left. (E) The total numbers of locally irrational topologies observed in all initial 3D structures (three structures for each RNA) and the predicted 3D models by IsRNAcirc (five models for each RNA, if available) were displayed in a type-dependent manner. All 34 circular RNAs were considered.

Notably, our IsRNAcirc can fix most of these aforementioned locally irrational topologies observed in the initial 3D structures through force field-guided simulations. As shown in Figs 3E and S3, the numbers of observed template-free loops (from 54 to 0), backbone tangles (between 80 and 28), base collisions (from 40 to 15), and sharp twists (from 62 to 60) are all largely decreased in the 3D models predicted by IsRNAcirc. It is worth noting that the number of initial 3D structures per RNA molecule is three, but the number of predicted 3D models by IsRNAcirc for each circular RNA is five. Therefore, the occurrence of these locally irrational topologies is significantly decreased in predicted 3D models (Fig S3). These results demonstrated that our IsRNAcirc is a very promising model for predicting circular RNA 3D structures, especially for those molecules with very long size.

### Representative predictions for each type of circular RNAs

To further illustrate the ability of IsRNAcirc in predicting circular RNA 3D structures, four representative examples were selected for each circular RNA type, and their prediction results were analyzed separately below.

For helix-circular RNAs, the POLR2A molecule consisting of 336 nucleotides was selected, and the prediction results are shown in Fig 4. According to the predicted 2D structure by cRNAsp12, the BSJ of POLR2A circular RNA is located in a 6-bp stem (Fig 4A). The initial linear 3D structure and the predicted 3D model by IsRNAcirc adopt different global folds, and the RMSD between these two structures is 20.7 Å (Fig 4B). Most importantly, the total number of locally irrational topologies of this molecule has decreased from 3 template-free loops (over all three initial linear structures) to 0 (over all five models predicted by IsRNAcirc). For example, a template-free loop of 14-nucleotide length (U120-A133) in the initial structure (caused by the lack of an appropriate template in the library, Fig 4C) was changed to a segment with a more reasonable geometry in the predicted model (Fig 4D). Furthermore, similar improvements in local topology were also observed after IsRNAcirc optimization for other helix-circular RNAs, such as ASXL1 (195 nucleotides) and KIAA0368 (435 nucleotides) molecules in Fig S3A.

**Figure 4.**
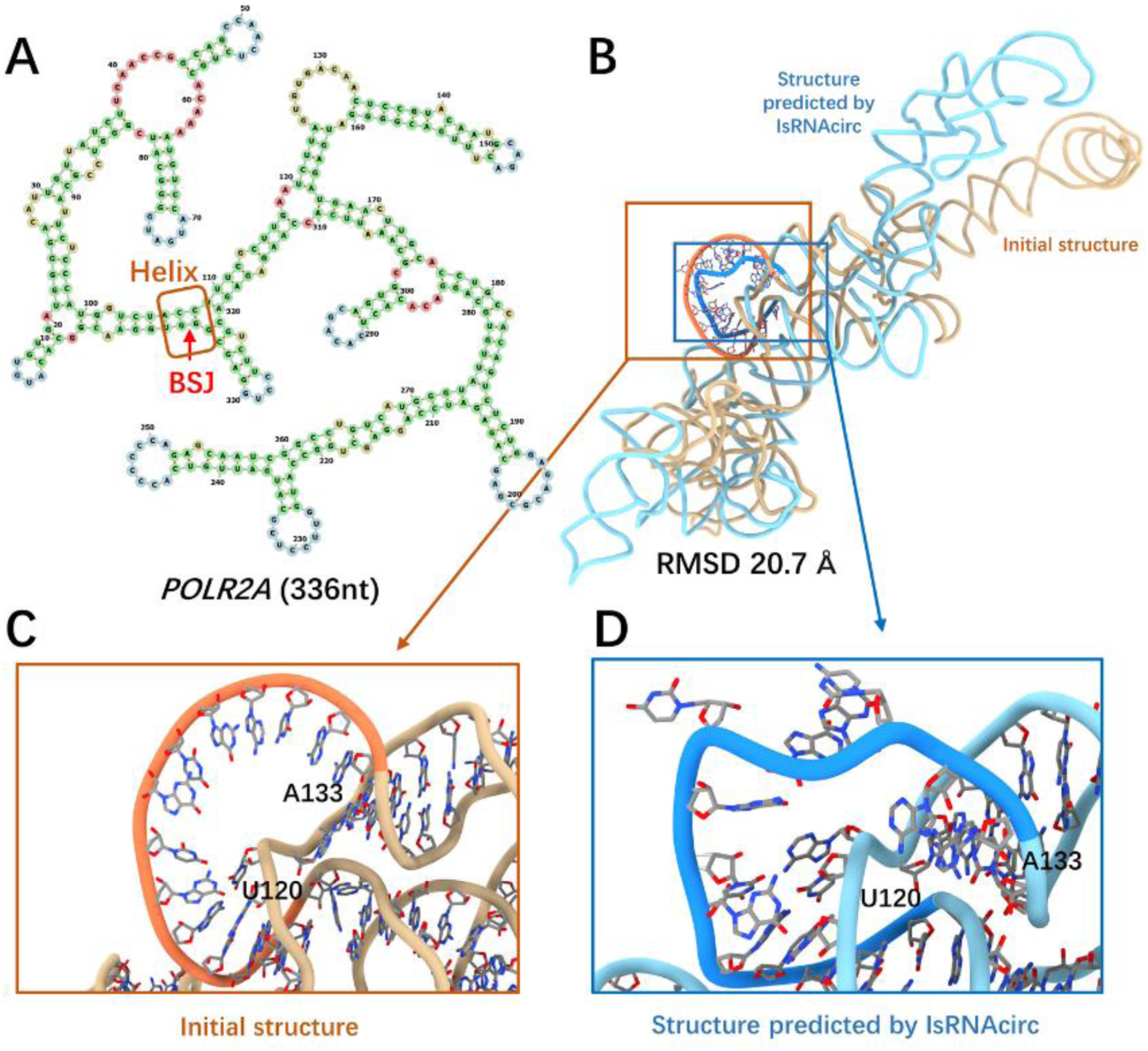
A representive prediction for helix-circular RNAs. (A) Predicted 2D structure of POLR2A consisting of 336 nucleotides (circRNA ID: hsa_circ_0000741), in which the BSJ is located in the helical region (marked by red arrow). (B) Superposition of the initial linear 3D structure (colored by light brown) and the predicted 3D structure of POLR2A circular RNA by IsRNAcirc (colored by light blue). (C-D) Zoom-in show the local details of 3D models: (C) the presence of template-free loop in the initial 3D structure and (D) the corresponding segment in the predicted 3D model by IsRNAcirc. The atoms carbon, oxygen, and nitrogen are colored by silver, red, and blue, respectively.

The FKBP8 molecule, composed of 259 nucleotides, was chosen as a representative case of hairpin-circular RNAs. Based on the predicted 2D structure shown in Fig 5A, the BSJ of this circular RNA is located in a 7-nucleotide hairpin loop. The RMSD between the initial linear structure and the predicted 3D model by IsRNAcirc is 27.7 Å (Fig 5B). As shown in Fig 5C, a backbone tangle between two segments (C25-U28 and A195-C200) was observed in the initial linear structure. After IsRNAcirc optimization, this backbone tangle was repaired in the predicted 3D model (Fig 5D). Overall, the total number of observed locally irrational topologies for the FKBP8 circular RNA decreases from 7 (over all three initial linear structures) to 5 (over all five models predicted by IsRNAcirc). For other hairpin-circular RNAs, such as ASAP1 (229 nucleotides) and RELL1 (434 nucleotides), a notable reduction in the number of locally irrational topologies was also found in the 3D models that predicted by IsRNAcirc (Fig S3B).

**Figure 5.**
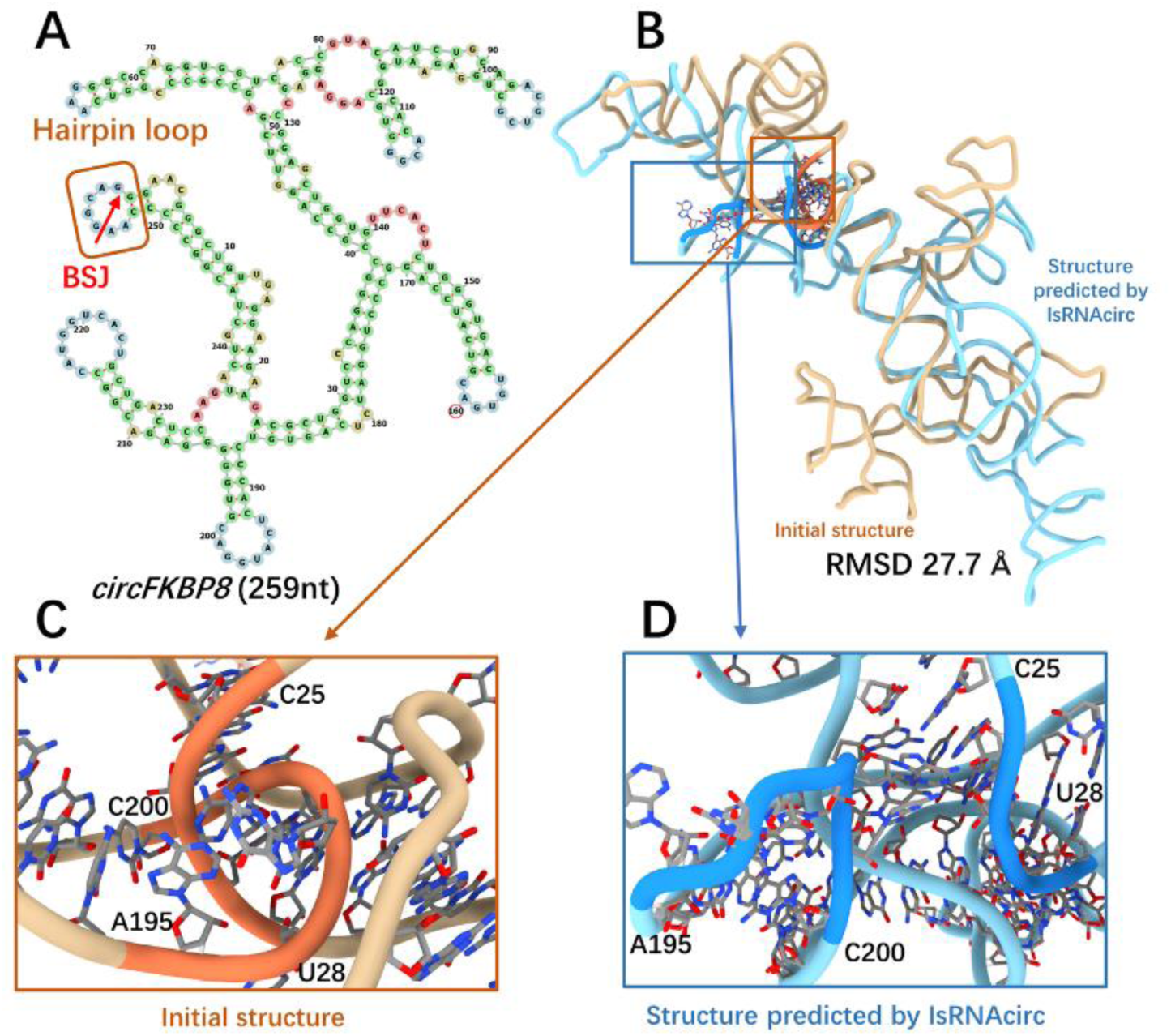
A representative prediction for hairpin-circular RNAs. (A) Predicted 2D structure of FKBP8 consisting of 259 nucleotides (circRNA ID: hsa_circ_0000915), in which the BSJ is located in the hairpin loop (marked by red arrow). (B) Superposition of the initial linear 3D structure (colored by light brown) and the predicted 3D structure of FKBP8 by IsRNAcirc (colored by light blue). (C-D) Zoom-in show the local details of 3D models: (C) the presence of a backbone tangle in the initial 3D structure and (D) the optimized segments in the 3D model predicted by IsRNAcirc. The atoms carbon, oxygen, and nitrogen are colored by silver, red, and blue, respectively.

For internal-circular RNAs, the MBOTA2 circular RNA composed of 224 nucleotides was selected as a representative example, and the prediction results are displayed in Fig. 6. Based on the 2D structure predicted by cRNAsp12, the BSJ of this molecule is located in a 1×1 internal loop (Fig 6A). The initial linear 3D structure and the predicted 3D model of the MBOTA2 molecule through IsRNAcirc adopt similar global folds in some regions, and the RMSD between these two structures is 19.5 Å (Fig 6B). Notably, no locally irrational topology was found in the 3D models predicted by IsRNAcirc, relative to 3 backbone tangles and 6 sharp twists observed in the initial linear structures. For instance, a sharp twist of the backbone segment (U164-G166) was found in the initial structure (Fig 6C), and this unreasonable topology was fixed by force field-guided MD simulations in IsRNAcirc (Fig 6D). In addition, similar repairs of locally irrational topologies by simulations in IsRNAcirc were also observed in other internal-circular RNAs, such as CNNB1 (378 nucleotides) and PVT1 (410 nucleotides) circular RNAs (Fig S3C).

**Figure 6.**
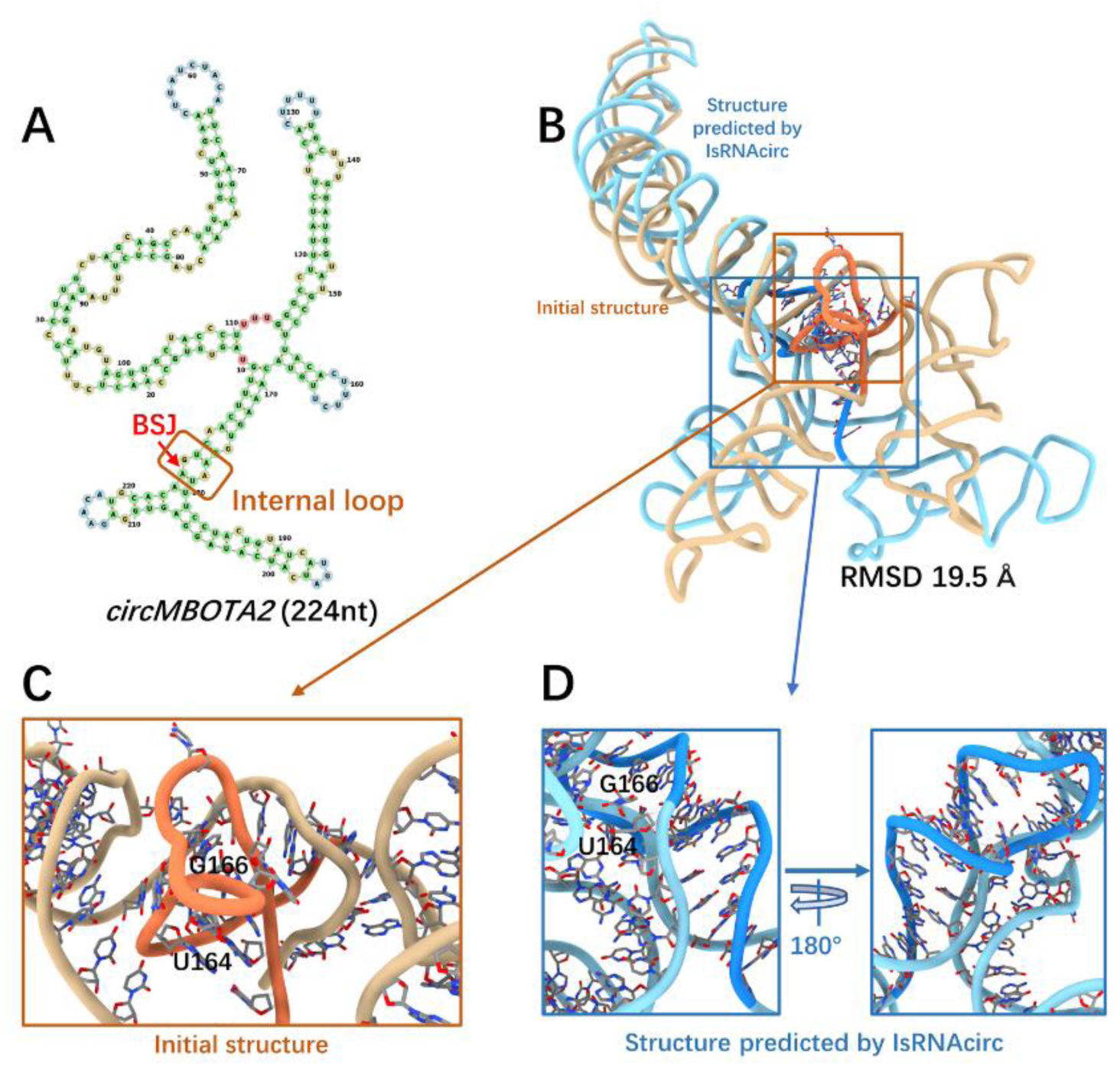
A representative prediction for internal-circular RNAs. (A) Predicted 2D structure of MBOTA2 consisting of 224 nucleotides (circRNA ID: hsa_circ_0007334), in which the BSJ is located in the 1×1 internal loop (marked by red arrow). (B) Superposition of the initial linear 3D structure (colored by light brown) and the predicted 3D structure of MBOTA2 by IsRNAcirc (colored by light blue). (C-D) Zoom-in show the local details of 3D models: (C) the presence of sharp twist of backbone in the initial 3D structure and (D) the optimized topology in the 3D model predicted by IsRNAcirc. The atoms carbon, oxygen, and nitrogen are colored by silver, red, and blue, respectively.

The SLC22A23 molecule, consisting of 259 nucleotides, was chosen as an illustrated example of junction-circular RNAs. According to the predicted 2D structure shown in Fig 7A, the BSJ of SLC22A23 circular RNA is located in a 3-way junction loop. Due to the great difference in the topological structure of multi-way junctions (circular RNA) and open-loops (linear counterparts), the initial linear structure and the predicted 3D model by IsRNAcirc adopt very different global folds, which results in a large RMSD value of 31.8 Å (Fig 7B). As shown in Fig 7C, there is a base collision in the local region of the initial 3D structure, where a backbone segment (U42-U46) passes through the middle space of two nucleobases (G16 and A17). After IsRNAcirc optimization, this irrational topology was fixed in the predicted 3D model, and a more reasonable conformation was presented. In total, the number of observed locally irrational topologies for the SLC22A23 molecule decreases from 3 (over all three initial linear structures) to 1 (over all five models predicted by IsRNAcirc). Moreover, for other junction-circular RNAs, such as EPHB4 (362 nucleotides) and PTK2 (394 nucleotides), the number of observed locally irrational topologies is also significantly decreased (Fig S3D).

**Figure 7.**
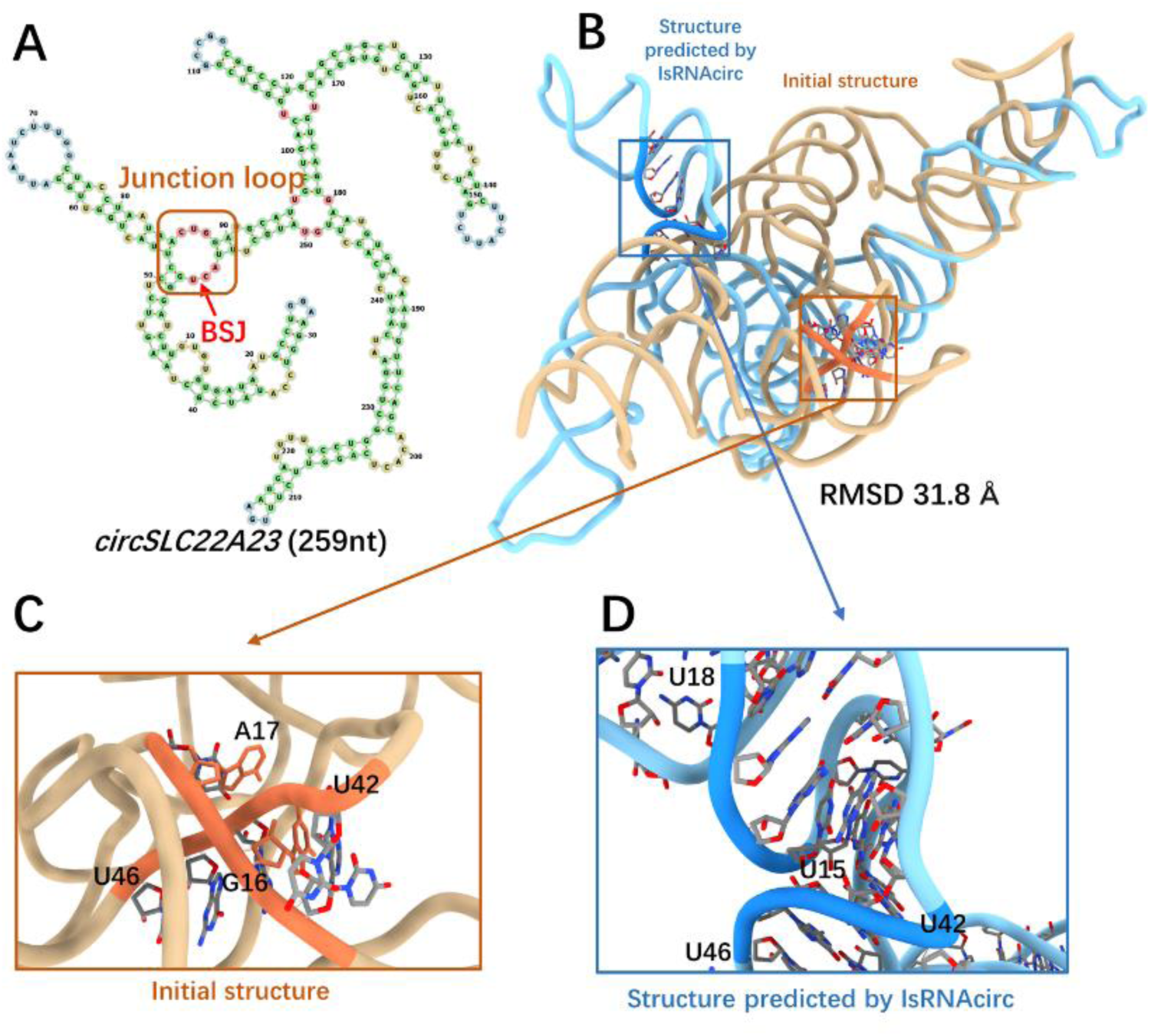
A representative prediction for junction-circular RNAs. (A) Predicted 2D structure of SLC22A23 consisting of 259 nucleotides (circRNA ID: hsa_circ_0075504), in which the BSJ is located in the 3-way junction loop (marked by red arrow). (B) Superposition of the initial linear 3D structure (colored by light brown) and the predicted 3D structure of SLC22A23 by IsRNAcirc (colored by light blue). (C-D) Zoom-in show the local details of 3D models: (C) the presence of base collision in the initial 3D structure and (D) the corresponding segments in the 3D model predicted by IsRNAcirc. The atoms carbon, oxygen, and nitrogen are colored by silver, red, and blue, respectively.

### Comparison with 3dRNA in circular RNA 3D structure prediction

Due to the current absence of experimental 3D structures of circular RNAs, the performance of IsRNAcirc in circular RNA 3D structure prediction cannot be directly validated or compared with the template-based 3dRNA approach. Therefore, apart from the aforementioned plausibility examination of the local topologies, an indirect comparison was also performed based on scoring functions. Previously, various scoring functions (energy functions) have been developed to select the near-native conformations from decoy structures, including 3dRNAscore[33], RASP[34], rsRNASP[35], DFIRE-RNA[36], and so on. In general, due to the energy function nature of most scoring functions, conformations with lower scores are considered closer to the native state. Consequently, an approach that can generate low-scoring conformations is better than one that can only obtain high-scoring structures. Here, we employed two knowledge-based scoring functions, DFIRE-RNA[36] and rsRNASP[35], to assess the predicted 3D structure models generated by IsRNAcirc and 3dRNA. To be fair, the same sequence and 2D structure inputs were used for both IsRNAcirc and 3dRNA, and the predicted 3D models were scored individually by those two scoring functions. As shown in Fig 8, based on the DFIRE-RNA scoring scheme, our IsRNAcirc can always generate lower energy conformations than the 3dRNA method for majority of tested circular RNAs with different circularized types. For example, for KIAA0368 (helix-circular RNA, 435 nucleotides) and EPHB4 (junction-circular RNA, 362 nucleotides) molecules, the average scores (-3.37×10^5^ and -2.80×10^5^ e.u.) of 3D models predicted by IsRNAcirc are obviously lower than the average scores (-3.01×10^5^ and -2.41×10^5^ e.u.) of models generated by 3dRNA. Furthermore, consistent results were also observed based on the alternative rsRNASP scoring function (Fig S4). Overall, these results indirectly demonstrate that IsRNAcirc has superior performance in circular RNA 3D structure prediction compared to the templated-based 3dRNA method, which mainly benefits from the more powerful sampling capability throughout the conformational space and accurate energy functions involved in the *ab initio* prediction method (e.g., IsRNAcirc here).

**Figure 8.**
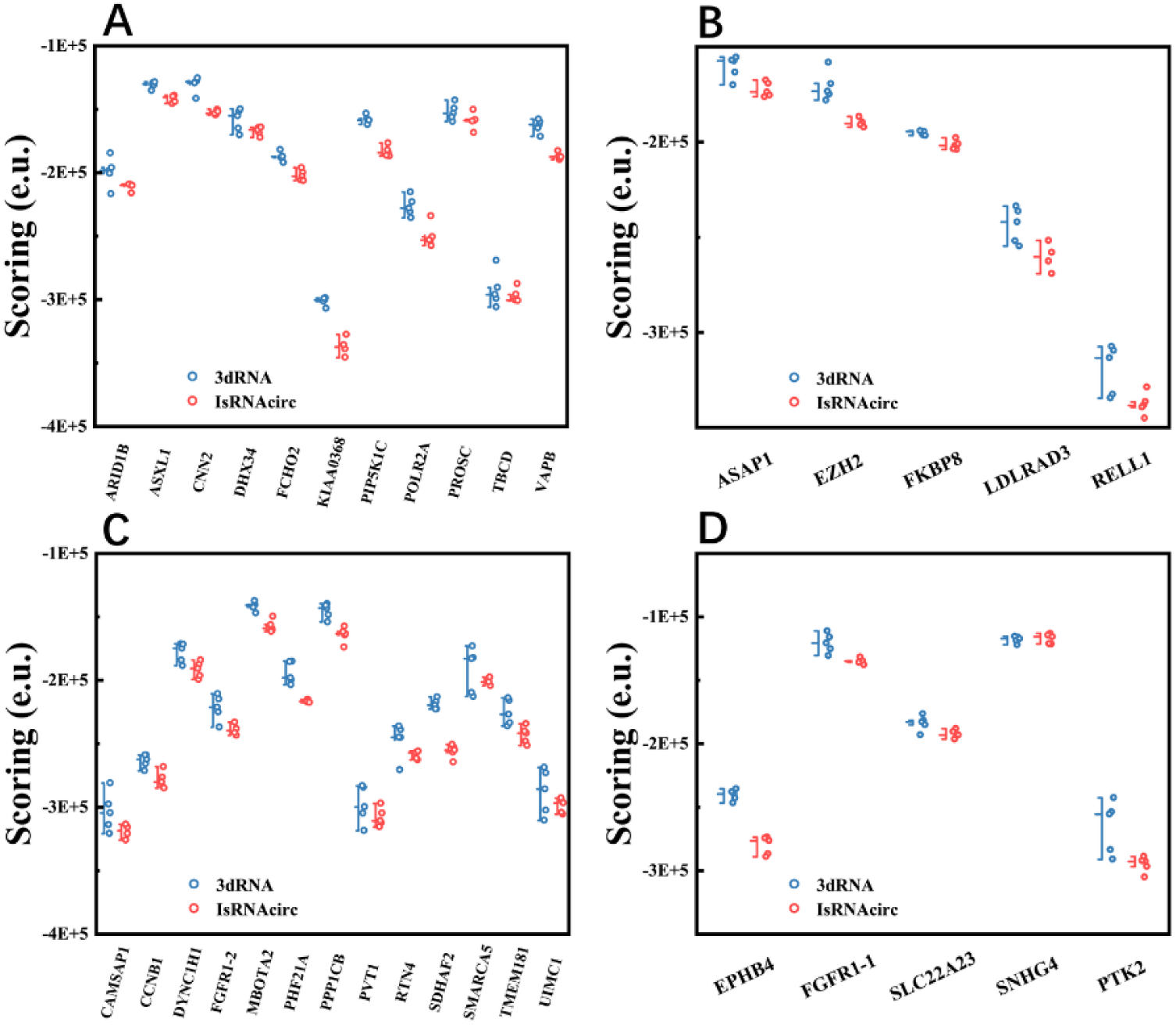
Comparison of the performance of circular RNA 3D structure predictions between IsRNAcirc and 3dRNA based on the DFIRE-RNA scoring function. Each 3D model predicted by IsRNAcirc or 3dRNA method was scored individually and displayed according to the circularized types: (A) helix-circular RNAs, (B) hairpin-circular RNAs, (C) internal-circular RNAs, and (D) junction-circular RNAs. e.u. (energy unit).

### Runtime

Runtime is one of the pivotal criteria for assessing the applicability and efficiency of any computational tool. Because IsRNAcirc relies on MD simulations to sample the huge conformational space, the running time consumed by IsRNAcirc to predict the 3D structures of circular RNAs is relatively long. For instance, IsRNAcirc took about 1,593 total CPU hours (Intel(R) Xeon(R) CPU E5-2620 v4 @ 2.10GHz) to generate the predicted 3D models for PIP5K1C molecule (consisting of 249 nucleotides). In addition, the relationship between runtime of IsRNAcirc and the length of circular RNA was also analyzed and the result was displayed in Fig 9. Notably, the total CPU time for predicting circular RNA 3D structure using IsRNAcirc increases almost linearly with the length of RNA molecules, mainly due to the coarse-grained nature of IsRNAcirc method. To further enhance the usability and practicality of IsRNAcirc in predicting long-size circular RNA 3D structures, several approaches will be considered in the future to improve computational efficiency, such as optimizing the source code, utilizing parallel computing technique, and introducing graphics processing unit (GPU)-based acceleration.

**Figure 9.**
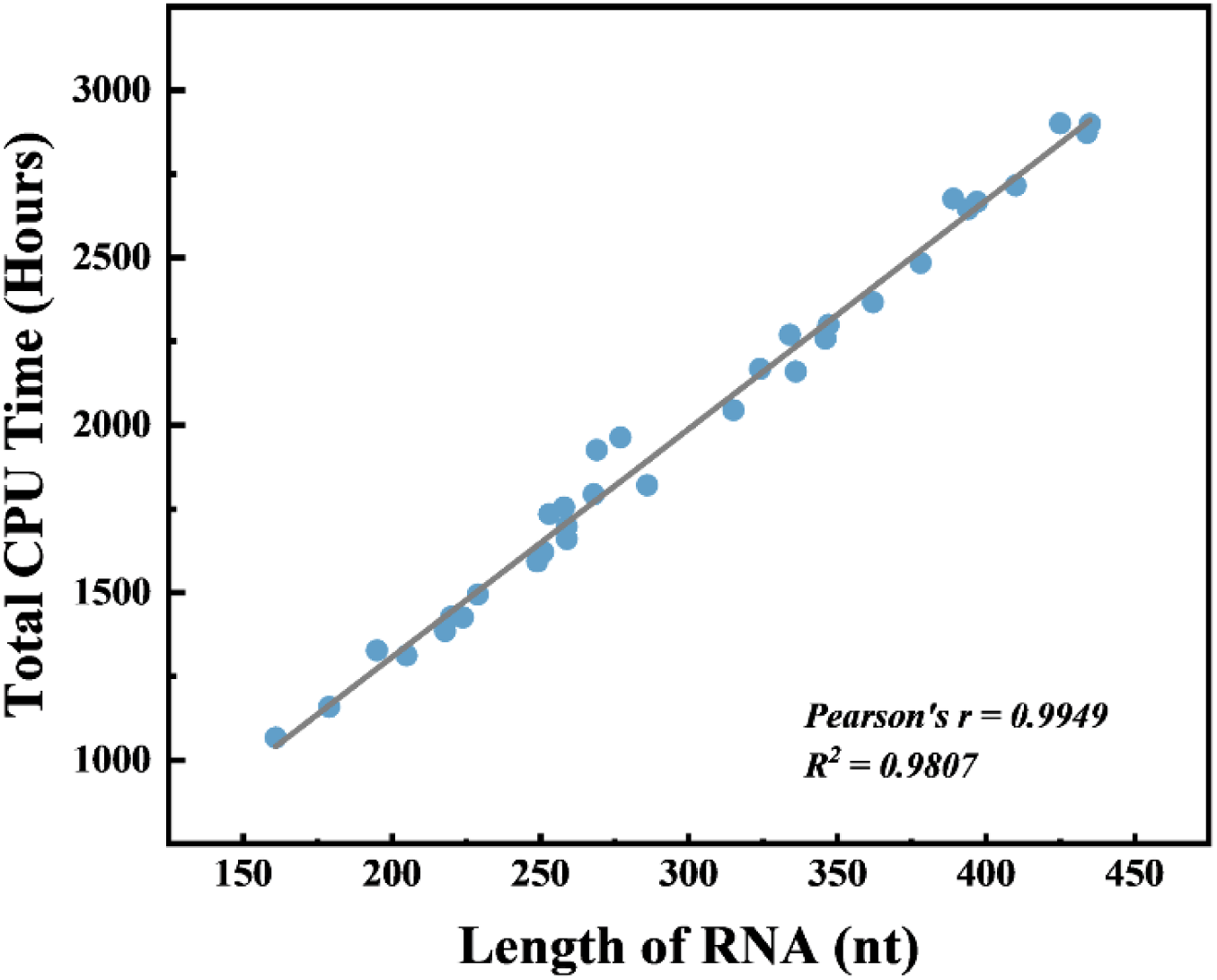
Runtime (total CPU time) in IsRNAcirc as a function of length of circular RNAs. Circles represent 34 circular RNAs and the line indicates the fitting result. Predictions were performed on intel(R) Xeon(X) CPU E5-2620 v4 @ 2.10GHz.

## Conclusion

In summary, based on our previous IsRNA2 method for linear RNA 3D structure prediction, we have developed an extended IsRNAcirc model to enable 3D structure prediction of circular RNAs. Based on the CG representation of nucleotides and the derived force field, IsRNAcirc employs REMD simulations to sample the huge conformational space and generate the most probable structures as the predicted 3D models of circular RNAs. Compared with the initial 3D structure of the linear counterpart obtained from RNAComposer, the local topologies of circular RNA 3D models predicted by IsRNAcirc show significant improvements. Although the performance of IsRNAcirc in predicting circular RNA 3D structures cannot be directly assessed at present due to the lack of experimental 3D structures, an indirect evaluation was performed utilizing energy-based score functions and then compared with the template-based 3dRNA method. For a test set comprising 34 circular RNAs (spanning in size from 161 to 435 nucleotides), our IsRNAcirc can generate 3D models with lower scores than 3dRNA in all the cases, regardless of the circularized type of circular RNAs. These results demonstrate that the proposed IsRNAcirc is a promising method to explore the structural details along with intricate interactions of circular RNAs.

## Methods

### Simulation details

All MD simulations were implemented in the LAMMPS[37] software with modified source code, and the Langevin dynamics (NVT ensemble) with an integration timestep Δ*t* = 1*fs* was performed. During the end closure stage (closing the 5’- and 3’-end of circular RNAs), a harmonic potential of equilibrium length *b*_0_ = 3.8 Å and a gradually increasing force constant (from *k* = 0.001 to 5.0 *kcal*/*mol*/Å^2^) were imposed on bead P of the first nucleotide and bead S of the last nucleotide. Simulated annealing simulations (from temperature T = 500 to 300 K) with a length of 10-50 ns (depending on the separated distance between the two ends) were run to sufficiently relax the initial structures and avoid unphysical structures. Meanwhile, to reduce the locally irrational elements involved in the initial structures, weakened base-pairing interactions (multiplied by a factor of 0.01/0.1/1.0) were also used during the simulated annealing simulations. For each initial structure, the most probable conformation (identified by clustering method) in the last 10% MD trajectory was selected as the starting structure for the subsequent simulation. During the structure prediction stage, REMD simulations with 10 replicas with temperatures ranging from 280 to 460 K were performed to efficiently sample the 3D conformational space. The simulation time for each replica is 100 ns, and three duplicated runs with different starting structures (if available) were run. Here, standard energy parameters were used for the relevant bonds/bond angles/torsion angles between two end nucleotides. After sufficient relaxation, the structure snapshots were collected from the last 50 ns simulations in the interval of 100 ps to construct the conformational ensemble (15,000 structures in total). Then, following the same steps as in IsRNA2, the top 10% structures with the lowest potential energies were submitted to the clustering procedure to generate the 3D predictions.

### Input and output

IsRNAcirc requires the sequence information, 2D structure (in dot-bracket format), and an initial 3D structure (in PDB format) of the corresponding linear RNA as input to predict circular RNA 3D structures. From the sequence information, the 2D structure of circular RNAs was predicted by the cRNAsp12 tool[31] (http://xxulab.org.cn/crnasp12/) using the default set. As multiple 2D structure predictions were provided by cRNAsp12, we selected the structure with the lowest folding free energy for further analysis. Using the sequence and predicted 2D structure as input, RNAComposer[18] (https://rnacomposer.cs.put.poznan.pl/) was utilized to obtain the initial 3D structures (three models, if available) of the corresponding linear RNAs. Additionally, another input of IsRNAcirc is a configuration file that specifies relevant parameters of the prediction process, which includes the simulation steps (simulation time), output parameters for collection of the conformational ensemble, cutoff threshold for clustering, the desired number of predicted models, and other relevant parameters.

The primary output of IsRNAcirc is the predicted 3D models (in PDB format) for given circular RNAs. By default, five 3D models are generated, which are the centroid structures of the top five largest clusters (if available) and then recovered to all-atom models through a built-in single-nucleotide fragment match algorithm[26]. In addition, users can flexibly obtain different numbers of predicted 3D models by modifying the relevant output parameter in the configuration file. Optionally, QRNAS[32] was employed to refine the predicted models by fixing possible errors in local geometries (such as unphysical bond lengths) and reducing the clash scores between atoms. Moreover, other useful outputs include simulation trajectories and corresponding energy files, log files of the clustering procedure, *etc*.

### Test dataset collection

To evaluate the predictive power of IsRNAcirc, a test dataset was collected from the previous study[30]. This test dataset consists of 34 circular RNAs with varying lengths ranging from 161 to 435 nucleotides, as listed in Supplementary Table 1. According to the location of BSJ in the 2D structure, these 34 circular RNAs can be categorized into four groups: 11 helix-circular RNAs, 5 hairpin-circular RNAs, 13 internal-circular RNAs, and 5 junction-circular RNAs.

## Data and code availability

The source code of IsRNAcirc and the data present in this study are available at https://github.com/DongZhangRNA/IsRNAcirc.

## Supporting information

**S1 Text. Supplementary figures.**

(PDF)

**S1 Dataset. Summary of 34 circular RNAs in the test set.**

(XLSX)

## Acknowledgements

We thank the anonymous reviewers for their insightful comments during the peer review process.

## Author Contributions

**Conceptualization:** Dong Zhang, Ruhong Zhou, Yunguang Tong.

**Data curation:** Haolin Jiang, Yulian Xu.

**Formal analysis:** Haolin Jiang, Yulian Xu, Dong Zhang.

**Investigation:** Haolin Jiang, Yulian Xu.

**Methodology:** Haolin Jiang, Dong Zhang.

**Project administration:** Dong Zhang.

**Resources:** Haolin Jiang, Yulian Xu.

**Software:** Dong Zhang, Haolin Jiang.

**Supervision:** Dong Zhang, Ruhong Zhou, Yunguang Tong.

**Visualization:** Haolin Jiang, Yulian Xu.

**Writing – original draft:** Haolin Jiang, Yulian Xu.

**Writing – review & editing:** Haolin Jiang, Yulian Xu, Yunguang Tong, Ruhong Zhou, Dong Zhang.

